# Dissecting the role of Amerindian genetic ancestry and *ApoE* ε4 allele on Alzheimer disease in an admixed Peruvian population

**DOI:** 10.1101/2020.03.10.985846

**Authors:** M. Cornejo-Olivas, F. Rajabli, V. Marca, P.G. Whitehead, N. Hofmann, O. Ortega, K. Milla-Neyra, D. Veliz-Otani, N. Custodio, R. Montesinos, S. Castro-Suarez, M. Meza, L.D. Adams, P.R. Mena, R. Isasi, M.L. Cuccaro, J.M. Vance, G.W. Beecham, P. Mazzetti, M.A. Pericak-Vance

## Abstract

Alzheimer disease (AD) is the leading cause of dementia in the elderly and occurs in all ethnic and racial groups. *ApoE ε4* is the most significant genetic risk factor for late-onset AD and shows the strongest effect among East Asian populations followed by non-Hispanic White populations and has a relatively lower effect in African descent populations. Admixture analysis in the African American and Puerto Rican populations showed that the variation in *ε4* risk is correlated with the genetic ancestral background local to the *ApoE* gene. Native American populations are substantially underrepresented in AD genetic studies. The Peruvian population with up to ∼80 of Amerindian ancestry provides a unique opportunity to assess the role of Amerindian ancestry in Alzheimer disease. In this study we assess the effect of the *ApoE* ε4 allele on AD in the Peruvian population.

A total of 78 AD cases and 128 unrelated cognitive healthy controls were included in the study. Genome-wide genotyping was performed using the Illumina Global screening array. Global ancestry and local ancestry analyses were assessed. The effect of the *ApoE ε4* allele on Alzheimer disease was tested using a logistic regression model by adjusting for age, gender, and population substructure (first three principal components). Logistic regression results showed that *ApoE ε4* allele is significantly associated with AD in Peruvian population with the high-risk effect (OR = 5.02, CI: 2.3-12.5, p-value = 2e-4). The average values of the local ancestries surrounding the *ApoE* gene (chr19:44Mb-46Mb) have the highest proportion of Amerindian (60.6%), followed by European (33.9%) and African (5.5%) ancestral backgrounds.

Our results showed that the risk for AD from *ApoE ε4* in Peruvians is higher than we have observed in non-Hispanic White populations. Given the high admixture of Amerindian ancestry in the Peruvian population, it suggests that the Amerindian local ancestry is contributing to a strong risk for AD in *ApoE* ε4 carriers. Our data also support the findings of an interaction between the genetic risk allele *ApoE ε4* and the ancestral backgrounds located around the genomic region of *ApoE* gene.

## Background

Alzheimer disease (AD) is a neurodegenerative disease accounting for over 70% of dementia cases in individuals ≥70 years of age^1^. AD has a multifactorial etiology, with both genetic and non-genetic risk factors, with liability-scale heritability estimates based on twin studies ranging between 0.58 and 0.79 with over 25 genetic risk factors contributing to AS risk^2,3^.

The apolipoprotein E (*ApoE*) gene (19q13.32) is the strongest known genetic risk factor for AD explaining up to 6% of the liablity-scale phenotypic variance^4,5^. *ApoE* codes for a protein that transports cholesterol through the interaction with cell surface receptors^6^. There are three *ApoE* alleles, ε2, ε3 and ε4, defined by two polymorphisms rs429358 and rs7412, that code for three protein isoforms ApoE2 (Cys130, Cys176), ApoE3 (Cys130, Arg176) and ApoE4 (Arg130, Arg176)^7^.

The association of *ApoE* with AD risk differs between populations and is not clearly established in groups of Amerindian (AI) descent. The strongest association of *ApoE* and AD risk has been observed in East Asian (EA) populations (ε3/ε4 odds ratio OR: 3.1–5.6; ε4/ε4 OR: 11.8– 33.1) followed by non-Hispanic White (NHW) populations (ε3/ε4 OR: 3.2; ε4/ε4 OR: 14.9) ^8,9^. Its effect is weaker in African-descent and Hispanic populations (ε3/ε4 OR:1.1–2.2; ε4/ε4 OR: 2.2–5.7)^10-14^.Genetic studies examining the interaction of genetic ancestry and risk effect of the *ApoE* in Caribbean Hispanic populations (Puerto Rican and Dominican Republic) showed that the effect of the ε4 is correlated with the ancestral background around *ApoE* with the attenuated effect on African-originated haplotypes^15,16^.

Peruvian population exhibits ∼83% Amerindian ancestral background, higher than other Latin American populations, such as Mexico (50%), Chile (40%), Colombia (28%), Argentina (28%) and Puerto Rico (16%)^17-19^. Peruvian Native American inhabitants show ancestry of three ancestral groups that originated by the split of an ancient group that migrated down the Americas after diverging from the East Asians and crossing the Bering Strait^20^. By admixing together and with non-Native inhabitants that arrived after Peru’s Spanish colonization, these AI groups gave rise to the current Peruvian mestizo population, resulting from admixture with European (EU), Asian and small African (AF) component^18-20^. Distribution of *ApoE* alleles in a sample of Mestizo Peruvian population from northern Lima suggest large contribution of *ApoE* ε3 genotypes, with approximately 9.5 % of cases harboring one or two *ApoE ε4* alleles^21^. No previous published studies have been addresses association of *ApoE* and AD in Peruvian population.

The heterogeneous ancestral make-up of Peruvians provides a unique opportunity to study the effect of global and local Amerindian ancestry on the effect of the ε4 allele over the risk of AD. If the discrepancies seen in the effect of the *ε4* allele across populations is caused by social factors, global, but not local to *ApoE*, ancestry is expected to also be associated to AD risk. Our goal is to use data from the Peruvian population to assess the role of AI genetic ancestry and the *ApoE* gene on AD.

## Methods

### Study samples and ascertainment

Unrelated cases and controls were ascertained from the Instituto Nacional de Ciencias Neurologicas in Peru as part of a genetics study in AD. All cases were assessed by trained neurologists following NINCDS-ADRDA criteria for probable AD ^22^. Cognitively intact controls were screened using the Clock drawing test, and the Pfeffer functional activities questionnaire ^23,24^. Controls were defined as individuals with no evidence of cognitive problems and age of exam (AOE) higher than 65 years of age. The dataset contained 79 AD cases (67.0% female, mean age at onset (AAO) = 72.3 years [SD=8.4]) and 128 cognitively healthy controls (59.1 % female, mean AOE = 75.0 years [SD =6.6]). This study was approved by the Ethical Committee of Instituto Nacional de Ciencias Neurologicas of Lima and the IRB of the University of Miami. Miller School of Medicine.

### Genotyping and quality control procedures

Genome-wide genotyping have been performed using llumina Global Screening Array. Quality control (QC) analyses were performed using software PLINK v.2^25^. Variants with the call score less than 95%, minor allele frequency less than 0.01, or not in Hardy-Weinberg equilibrium (HWE) (p<1.e-6) were eliminated. The concordance between reported sex and genotype-inferred sex was checked using X-chromosome data. The relatedness among the individuals were assessed using “identical by descent” (IBD) allele sharing. *ApoE* genotyping was performed as in Saunders et al.^26^

### Assessment of Genetic Ancestry

Global ancestry was evaluated using GENESIS software program that is robust to known and cryptic relatedness^27^. Firstly, the KING-Robust kinship coefficient estimator was used to calculate the KING matrix that includes pairwise relatedness and measures of pairwise ancestry divergence^28^. PC-AiR method was then applied to calculate “preliminary” principal components (PC) by using KING matrix. Default kinship and divergence threshold values have been used. The PC-Relate method that uses “preliminary” PCs to account for the samples ancestry variation and calculate the genetic relationship matrix (GRM) that is robust for the population structure, admixture, and departure from HWE was applied to the data. PC-AiR method was once more applied to the data by using the robust kinship estimates (GRM) and calculated PCs that accurately capture population structure. PCs were calculated with and without population reference datasets. Four reference populations were used including AI, EU, AF, and EA from Human Genome Diversity Project (HGDP) data for the reference populations^29^.

To estimate the admixture proportion, a model-based clustering algorithm was performed as implemented in the ADMIXTURE software^30^. Supervised ADMIXTURE analysis was used at K = 4 by including the same four reference populations from HGDP reference panel we used in PC-AiR approach.

To assess the local ancestry, HGDP reference panel was combined with the Peruvian data using the PLINK v2 software including approximately the same number of individuals from three reference populations EU, AF and AI (∼ 100). Then, all individuals in combined dataset were phased using the SHAPEIT tool ver. 2 with default settings and 1000 Genomes Phase 3 reference panel^31,32^. Finally, RFMix was performed using the discriminative modeling approach, to infer the local ancestry at each loci across the genome. We ran RFMix with the PopPhased option and a minimum node size of 5^33^.

The heterogenous risk effect of the *ApoE* gene across the populations is suggested to be correlated with the ancestral background local to the *ApoE* gene. Thus, to examine the ancestral background in our dataset we calculated the average ancestry proportions at the *ApoE* by taking the average of the local ancestry estimates around the *ApoE* gene (from 44 Mb to 46 Mb on chromosome 19)^15^. The pipeline to calculate the global and local ancestries was developed by our group using R and Python scripts.

### Statistical Analysis

To assess the effect of the *ApoE ε4* allele in Peruvian population we performed logistic regression approach. In this model, the association was tested between the affection status and gene dose of the *ApoE ε4* allele by adjusting for age, gender, and populations substructure (PC1, PC2, and PC3). Statistical analysis was performed using the “GLM2” package available in R computing environment^34^.

## Results

The supervised ADMIXTURE analysis showed that Peruvians are four-way admixed population with the 63.6% AI, 29.7% EU, 3.8% AF and 2.9% EA ancestral background. Figure 1A shows the box-plot of the average ancestry across the all individuals in the dataset. The ancestral proportion of each individual is illustrated in the bar-plot Figure 1B, where each column reflects the admixture structure of a single individual as the proportion of different colors. Admixture analysis results confirm the recent genetic studies showing a four-way admixture (AI, EU, AF, and EA) structure in Peruvians.

**Figure 1.**
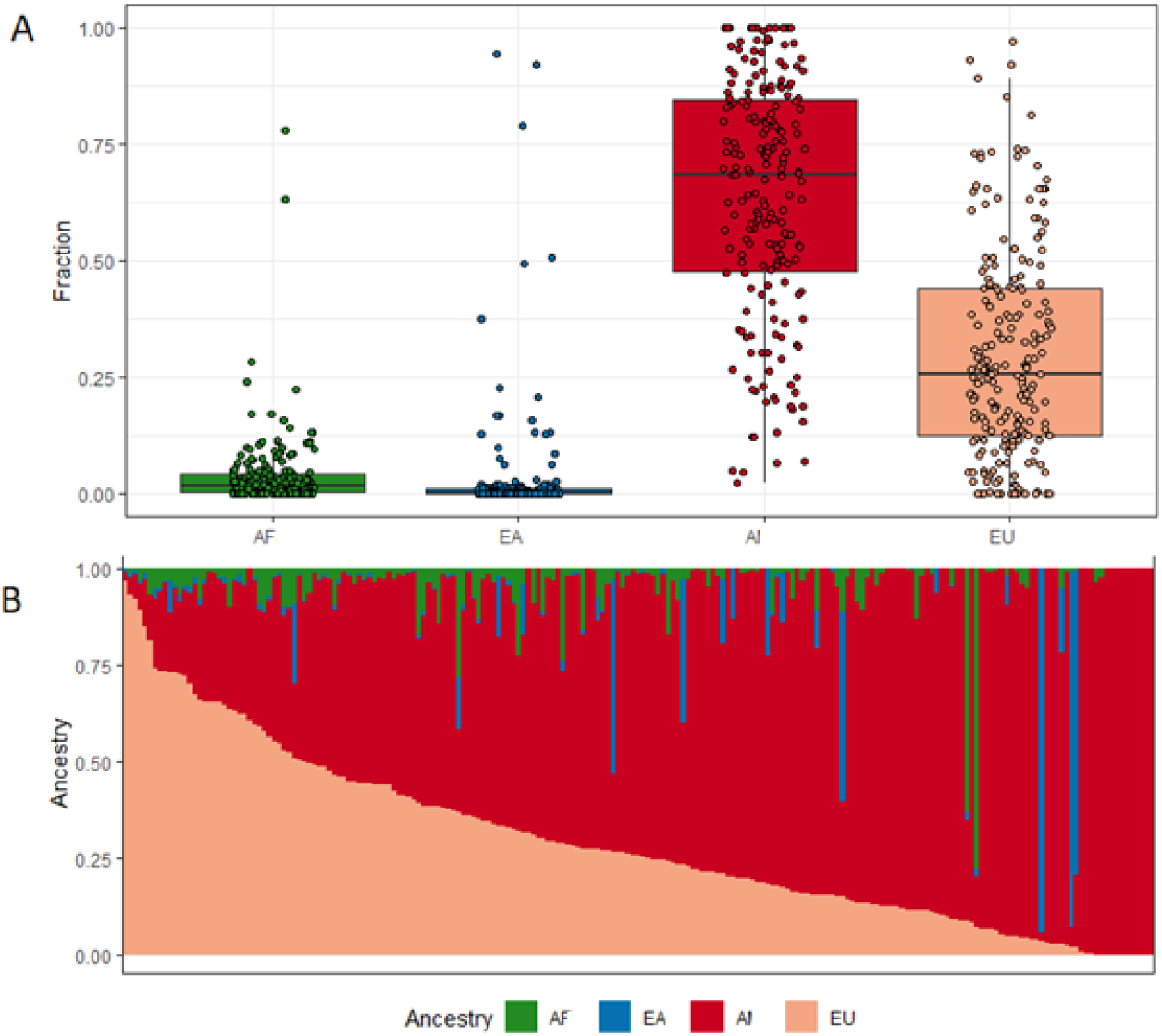
A. The box-plot of the four parental ancestries in Peruvian dataset B. Bar-plot of four-way admixed Peruvian individuals estimated using ADMIXTURE software at K = 4

The allele frequency distribution of the *ApoE* alleles are illustrated in Table 1. The affected individuals have higher frequency of *ApoE ε4* allele (9.2%) than individuals (4.6%) that are cognitively normal. Logistic regression results showed that the *ApoE ε4* allele is significantly associated with AD in Peruvian population with the high-risk effect (OR = 5.02, CI: 2.3-12.5, p-value = 2e-4). The average of the local ancestries around the *ApoE* gene showed that the distribution of the parental ancestries local to the *ApoE* gene is the similar to the average ancestry across the genome with the highest proportion of AI (60.6%), followed by EU (33.9%) and AF (5.5%) ancestral backgrounds.

**Table 1.** ApoE genotype and allele frequencies in cases and controls.

|  | Cases (%) | Controls (%) |
| --- | --- | --- |
|  | *Genotypes* |  |
| $\epsilon 2\epsilon 3$ | 3(1.4) | 8(3.9) |
| $\epsilon 3\epsilon 3$ | 43(20.8) | 102(49.3) |
| $\epsilon 3\epsilon 4$ | 28(13.5) | 17(8.2) |
| $\epsilon 4\epsilon 4$ | 5(2.4) | 1(0.5) |
| *Alleles* |  |  |
| $\epsilon 2$ | 3(0.7) | 8(1.9) |
| $\epsilon 3$ | 117(28.3) | 229(55.3) |
| $\epsilon 4$ | 38(9.2) | 19(4.6) |
| **Total** | **79** | **128** |

## Discussion

The *ApoE* ε4 allele is the most significant genetic risk factor for late-onset AD with the differences in effect size among the populations. Our results showed that the risk for AD from *ApoE ε4* allele in Peruvians is higher than we have observed in NHW populations. Given the high admixture of AI in the Peruvian population, it suggests that the AI local ancestry is contributing to a strong risk for AD in *ApoE* ε4 carriers. This would align with the current believed migration pattern of AI from East Asia, where *ApoE* ε4 carriers have the highest *ApoE* ε4 risk for AD.

The prevalence of AD varies among the diverse populations. Moreover, AD genetic studies in different ethnic groups have shown variation in both risk effect size and variants (e.g. *ApoE, ABCA7, SORL1, etc*.)^8,35-37^. This heterogeneity suggests that distinct genetic architecture can lead to differing disease susceptibility. Thus, studying the diverse populations is critical to the understanding of the molecular mechanism underlying the disease pathogenesis and the success of precision medicine. However, diverse populations and especially populations with the AI ancestry are substantially underrepresented in AD genetic studies. The Peruvian population with a large proportion of AI ancestry provides a unique opportunity to assess the role of AI ancestry in AD. This study by confirming the correlation of the genetic ancestry with the risk effect in ε4 allele shows the importance of studying different populations to evaluate the ancestry-specific genetic modifiers correlated with ancestry. Ultimately, studying diverse populations is essential to understand the genetic factors initiating AD pathogenesis that may contribute to health disparities and ultimately the development of effective therapies.

